# Differential biotransformation ability may alter fish biodiversity in polluted waters

**DOI:** 10.1101/2024.07.26.605280

**Authors:** Marco E. Franco, Juliane Hollender, Kristin Schirmer

**Affiliations:** Department of Environmental Toxicology, Swiss Federal Institute of Aquatic Science and Technology, Eawag, 8600 Dübendorf, Switzerland; Department of Environmental Chemistry, Swiss Federal Institute of Aquatic Science and Technology, Eawag, 8600 Dübendorf, Switzerland; Department of Environmental Systems Science, ETH Zürich, 8092 Zürich, Switzerland

**Keywords:** Biotransformation, Fish, Biodiversity, Chemical pollution

## Abstract

Divergence in the activity of biotransformation pathways could lead to species sensitivity differences to chemical stress. To explore this hypothesis, we evaluated the biotransformation capacity of five fish species that are representatives of Swiss biodiversity assemblages and that inhabit watercourses surrounded by different land use. We report important interspecific differences regarding the presence and activity of major biotransformation pathways, such as the invasive pumpinkseed (*Lepomis gibbosus*) displaying micropollutant clearance between 3- and 7–fold higher than native species (e.g. *Salmo trutta, Squalius cephalus*) collected in the same areas. These differences were exacerbated by urban and agricultural influence, which increased biotransformation potential at the enzyme level by as much as 11-fold and micropollutant clearance by approximately 2-fold compared to biotransformation levels in areas with minimal human influence. In the context of the chemical defensome, we argue that fish with low biotransformation activity carry a greater burden on chemical stress, making them less likely to cope with additional stressors and sustain their population in competition with species with a higher biotransformation capacity.

## 1. Introduction

The continuous introduction of legacy and novel chemicals into the environment severely compromises the biodiversity and ecological integrity of different ecosystems. This issue has become a matter of international concern among researchers and policymakers (*1, 2*). Aquatic ecosystems, in particular, are among the most threatened by pollution, as they represent important sinks for a large variety of chemicals (*3*). However, studies in echinoderms and in different fish species have indicated that organisms evolved an array of biological processes that act as defense mechanisms against chemical exposure, also referred to as the chemical defensome (*4, 5*). These evolutionary traits may allow some species to have a physiological advantage over others, unavoidably resulting in the decline of sensitive species inhabiting polluted environments. Consequently, generating knowledge about species-specific sensitivity to chemical pollution and its role in altering biodiversity represents one of the major challenges in ecotoxicology.

Among the different components of the chemical defensome, biotransformation pathways are imperative to maintain the fitness of exposed organisms; they facilitate chemical elimination from the body, thereby reducing bioaccumulation (*6*). Several biotransformation-associated genes are conserved across different taxonomic groups, likely due to the need of being equipped with defense mechanisms against endogenous and exogenous stressors (*5*).

However, their inducibility and translation into enzyme activity may differ across species. As a result, whether species are able to cope with chemical exposure not only depends on the sole presence of different biotransformation pathways but also on the species’ ability to display them, which in turn may be influenced by the exposure history of the populations and abiotic factors, such as water quality (*7, 8*).

Therefore, our hypothesis for this study was that biotransformation processes are suitable indicators of species sensitivity to pollution, assuming that species that display high biotransformation ability are, to some extent, more tolerant and therefore more abundant. We focused our efforts on wild freshwater fishes, representative of regional biodiversity assemblages, which also allowed us to account for the influence of habitat conditions, such as differential levels of pollution. We designed a comparative study in which enzymatic fractions were isolated from fish livers, characterized for biotransformation enzyme activity, and used to measure the clearance kinetics of six ubiquitously found micropollutants and their resulting biotransformation product (BTP) profiles.

## 2. Site selection and fish collections

We sampled six freshwater streams within the Aare catchment in Switzerland. The selected sites displayed different levels of anthropogenic activity and land use, according to a classification established for freshwater biodiversity monitoring campaigns in the Aare catchment by the Department of Fish Ecology and Evolution at Eawag (Fig. 1A, Table S1). A comparison of basic water quality parameters among collection sites indicated relatively similar temperature, pH, and dissolved oxygen (Fig. 1 B/C/D). However, a trend towards increasing specific conductivity was observed as the levels of anthropogenic activity became higher (Fig. 1E). Indeed, elevated conductivity has been directly associated with the presence of different chemicals in surface waters (*9, 10*), suggesting an important impact of the surrounding land use on the water quality of the sampled streams.

**Fig. 1.**
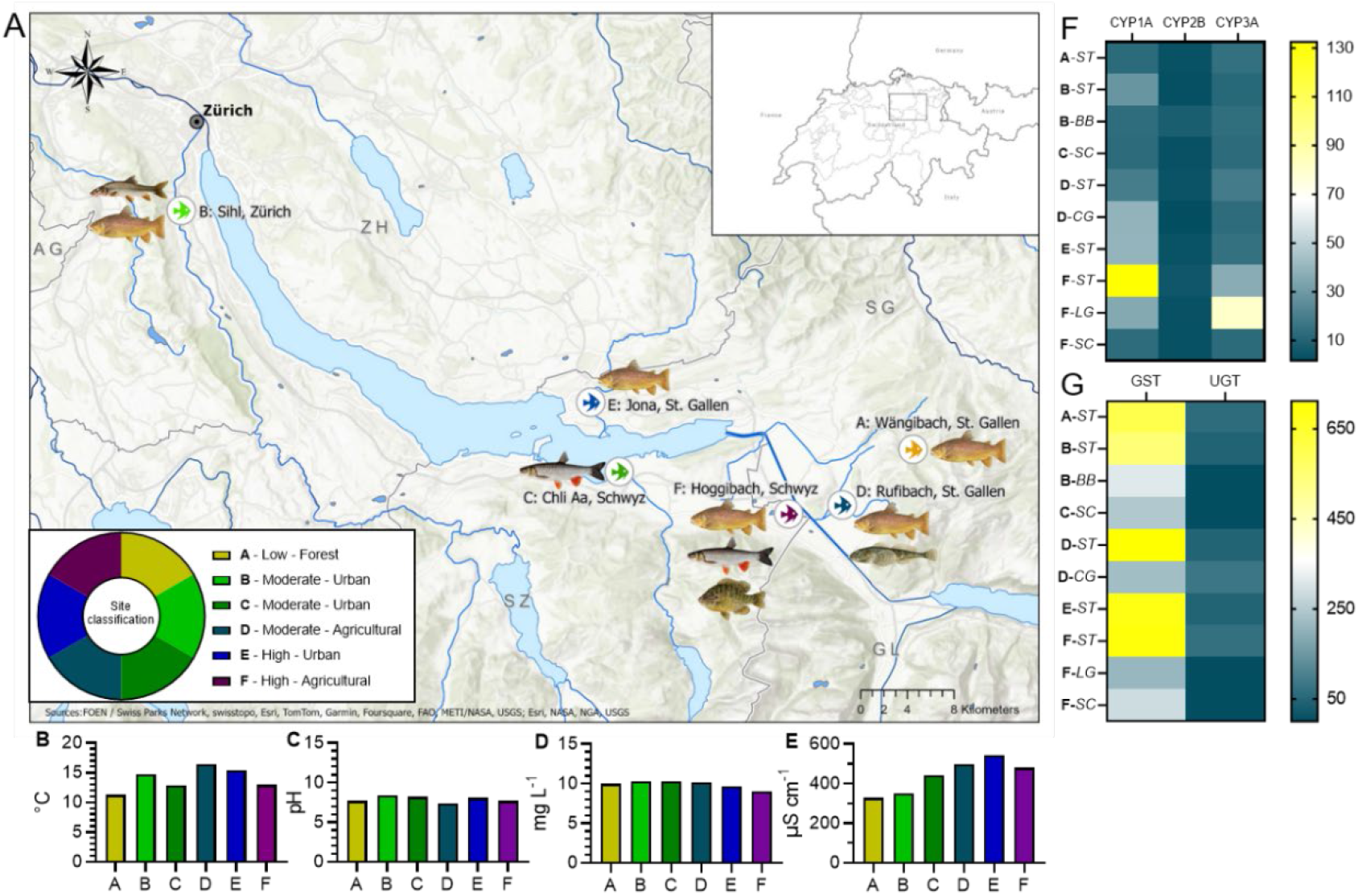
Physicochemical features of collection sites and biotransformation enzyme activity of five different fish species (*Salmo trutta* (ST), *Barbus barbus* (BB), *Cottus gobio* (CG), *Lepomis gibbosus* (LG), and *Squalius cephalus* (SC)) inhabiting surface waters across three cantons in Switzerland. Collection sites are organized according to the magnitude (low, moderate, high) and type (forest, urban, agricultural) of surrounding land use. Basic water parameters, including B) temperature, C) pH), D) dissolved oxygen, and E) specific conductivity are reported for each collection site. Further information about collection sites is shown in table S1. For all fish species, maximum hepatic activity rates (*Vmax*) were determined for F) phase I (CYP1A, CYP2B, CYP3A; pmol mg protein^-1^ min^-1^) and G) phase II (glutathione S-transferase (GST; nmol mg protein^-1^ min^-1^) and UDP-glucuronosyl transferase (UGT; pmol mg protein^-1^ min^-1^)) biotransformation enzymes. The displayed data for each biotransformation enzyme were derived from two S9 pools (one for *B. barbus*) for each species with two technical replicates.

Across all sites, we collected five fish species via non-lethal, backpack electrofishing: brown trout (*Salmo trutta*), chub (*Squalius cephalus*), pumpkinseed (*Lepomis gibbosus*), and the bottom-dwellers common barbel (*Barbus barbus*) and European bullhead (*Cottus gobio*). The presence of these fish species at the collection sites were in line with reports on fish biodiversity in Switzerland (*11*), which concluded that *S. trutta* is the most widespread species in Swiss surface waters (present in five of the six collection sites). Although invasive to European surface waters, *L. gibbosus* is also considered a representative species of regional biodiversity assemblages (*12*). Additionally, higher diversity occurred in lowland surface waters influenced by high anthropogenic activity (e.g. more species caught in sites influenced by agriculture) than in streams with low human influence. Fish were dissected on-site in accordance with animal experimentation regulations in Switzerland (Animal Testing Permit No. BE11/2022), and liver S9 sub-cellular (enzymatic) fractions were isolated following standard procedures (see *Isolation of liver S9 sub-cellular fractions* section in supplemental material; (*13, 14*))

## 3. Biotransformation enzyme activity differed among species and collection sites

S9 sub-cellular fractions were characterized for the presence and activity of major phase I and II biotransformation pathways (see *Biotransformation enzyme activity* section in supplemental material). Enzymes selected for phase I biotransformation were CYP1A, CYP2B, and CYP3A, given their important role in xenobiotic biotransformation in fish (*6*). At the same time, activity of these enzymes would reveal the presence of the aryl hydrocarbon receptor (AhR), the constitutive androstane receptor (CAR), and the pregnane X receptor (PXR) pathways, respectively (*15-17*). The enzymes selected for phase II were glutathione S-transferase (GST) and UDP-glucuronosyl transferase (UGT) given their direct role in supporting detoxification mechanisms across different fish species (*18-20*).

All of the species displayed detectable levels of the three phase I biotransformation enzymes (Fig. 1F and S1; Table S4), which followed Michaelis-Menten kinetics. This is particularly important because the target enzymes reached a level of saturation when substrate concentrations continued to increase. In other words, at specific levels of substrate chemicals biotransformation activity could not increase further (*21*). Among species, *S. trutta* displayed CYP1A activity between 3- and 11-times higher than the other species collected, whereas CYP2B- and CYP3A-like activities were higher in *B. barbus* and *L. gibbosus*, respectively. Furthermore, differences in CYP1A and CYP3A-like activities observed in *S. trutta* at the site with the highest human influence were more than 10- and 3-times higher, respectively, than in *S. trutta* from less polluted locations. Similar observations occurred for phase II biotransformation enzymes (Fig. 1G and S2; Table S4), as they displayed clear site- and species-specific trends, with *S. trutta* displaying GST activity more than 3-times higher than the other species. In addition, UGT activities were only detected in *S. trutta* and *C. gobio*, with the latter displaying the highest activity despite not showing Michaelis-Menten kinetics. This observation could potentially indicate a better efficiency of glucuronidation reactions despite high concentrations of substrate chemicals in *C. gobio*. Similar observations related to differential enzyme activity in fish were also recently highlighted for species of regulatory interest (*22*). In addition, different levels of human activity and habitat conditions that result in different inducibility could have also influenced differences in enzymatic activity.

To provide an comparison, previous studies focusing on enzyme activity and legacy pollutants (e.g. polycyclic aromatic hydrocarbons) have indicated habitat-related inducibility where the levels of pollution drove the activity of biotransformation enzymes in Atlantic killifish (*Fundulus heteroclitus*) and Gulf killifish (*Fundulus grandis*) populations inhabiting polluted waters (*23, 24*). These observations as well as the ones from our study indicating differential enzyme activity in fish illustrate the influence of habitat conditions on biotransformation pathways. In consequence, these habitat-driven biotransformation profiles could have important implications in reducing adverse effects when biotransformation ability is induced. However, one must also consider that such differences could also increase toxic outcomes when biotransformation activity remains low, as was the case for bottom-dwelling species.

We were further interested in whether enzymatic activity was translatable to the actual intrinsic clearance of omnipresent micropollutants in surface waters. This is because focusing solely on enzyme presence and activity across species may not fully reveal the true biotransformation ability of organisms. Previous studies have indicated that the molecular blueprint governing the affinity of biotransformation pathways towards different classes of pollutants may trigger variable biotransformation rates (*18, 25, 26*). For example, slight modifications in the ligand-binding domain of the AhR led to differential affinity towards toxic AhR agonists in birds. Such affinity differences caused reduced sensitivity towards the chemicals and a lower magnitude of adverse effects was observed for the species in which the AhR displayed weak binding to the compounds (*27*). Therefore, to further illustrate the role of biotransformation pathways in influencing across species sensitivity to pollution, we determined clearance rates (CL_*IN VITRO*_) of three pharmaceuticals: propranolol, diclofenac, paracetamol, and three pesticides: azoxystrobin, terbuthylazine, and pirimicarb, and evaluated the identity and formation rates of their biotransformation products (BTPs).

## 4. Fish species biotransform micropollutants at different rates

We detected significant species-specific clearance rates for four of the micropollutants: propranolol, diclofenac, azoxystrobin, and terbuthylazine (Fig. S3 A/B/D/E, Table S5). Biotransformation rates were significantly higher for propranolol and azoxystrobin (Fig. 2) than for diclofenac and terbuthylazine. No significant clearance was observed for paracetamol and pirimicarb (Fig. S3 C/F).

**Fig. 2.**
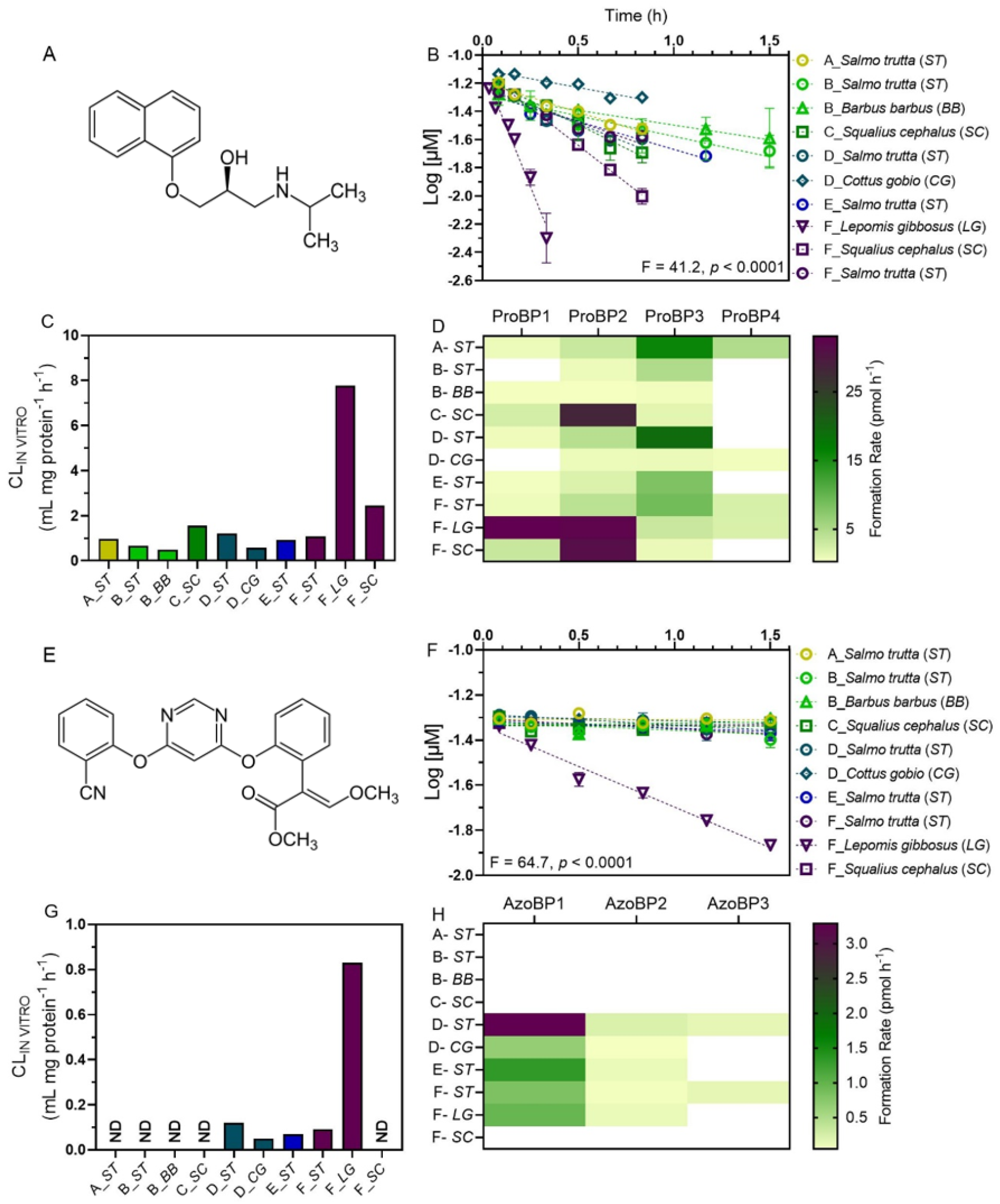
Hepatic biotransformation of A) propranolol and E) azoxystrobin in five different fish species inhabiting surface waters with different magnitude and type of surrounding land use. Collection sites are organized from light to dark colors as the magnitude of land use increased. Panels B and F illustrate substrate depletion experiments, that is the log-linear decrease of chemical concentrations over time (h). Data points correspond to the mean Log_10_ concentrations (µM) ± SEM of two independent experiments with each S9 pool (n = 4 for most species and n = 2 for *B. barbus*). Each graph indicates whether the slopes resulting from linear regression analyses are different from each other, when considering *p* < 0.05 as significant. Substrate depletion plots were then used to determine the first-order depletion rate constant (*k*_DEP_; 1/h), followed by the estimation of (C/G) hepatic intrinsic clearance (CL_IN VITRO_; mL mg protein^-1^ min^-1^). Panels D and H show the formation rates (pmol h^-1^) of different biotransformation products (BTPs), also calculated from hepatic biotransformation. Empty (white) cells indicate that a given BTP was not detected in the respective fish species.

Fish species collected at the location with high agricultural influence (site F) displayed between 2- and 16-times higher micropollutant clearance than fish collected from the other locations. In general, *L. gibbosus* displayed clearance 3-to 7-times higher than the other species, followed by *S. cephalus* and *S. trutta*, whereas bottom-dwelling species (*C. gobio* and *B. barbus*) displayed the lowest clearance. This same species-specific clearance profile was maintained across all the chemicals that were biotransformed. Although biotransformation studies with the invasive *L. gibbosus* are limited, previous studies with a close relative, *Lepomis macrochirus*, have indicated a high biotransformation potential of this species in comparison to cold-water species, like salmonids (*22*). Our observations indicate that this trend could be also true for *L. gibbosus* when compared to the native European species considered in our study. We also observed that the species that displayed high biotransformation potential at the enzyme activity level were able to clear the micropollutants faster (Table S4 and S5). For these particular compounds and in the context of the chemical defensome, it is likely that fishes with high biotransformation activity, like *L. gibbosus*, are able to cope with pollution and minimize toxic outcomes.

To illustrate this rationale, previous studies employing different fish species have indicated that environmental exposure to propranolol, the compound that had the highest biotransformation rates in our study, leads to abnormal heart rate and development (*28*). Such effects would be minimized when propranolol is efficiently excreted (*29*), even when aquatic organisms are continuously exposed in areas where propranolol is pseudo-persistent due to its ubiquitous presence in e.g. effluent-impacted waters. Based on human data, propranolol is mainly processed by CYP1A and CYP2D isoforms (*30*). Therefore, while the latter was not targeted in our study given the absence of a specific substrate and bioassay, it is likely that a similar isoform was of particular importance in *L. gibbosus* and, to a lower extent, in *S. cephalus*, given the high propranolol clearance in these species but low CYP1A activity in comparison to e.g. *S. trutta*. In this context, fish that are equipped with high biotransformation capacity could reduce the toxicological risks associated with chemicals for which biotransformation is an essential process for their elimination. In turn, such species with higher biotransformation ability would be better able to detoxify chemicals, maintain fitness, and outcompete sensitive species inhabiting contaminated waters.

## 5. Biotransformation product profiles followed species-specific patterns

The involvement of different biotransformation pathways in clearing propranolol across species was also evident from the formation of four biotransformation products (BTPs; Fig 2D; Table S6). Both ProBP1 and ProBP2, potentially corresponding to propranolol-N-hydroxyl and 4-hydroxypropranolol (*18, 31*), were favored in *L. gibbosus* and *S. cephalus*. Thus, it is possible that biotransformation activity by CYP isoforms similar to e.g. CYP2D6 (given its direct involvement in propranolol biotransformation in mammals) resulted in the formation of these two BTPs. Contrarily, ProBP3 may have been a primary BTP resulting from CYP1A-mediated biotransformation, as indicated by both the significantly high CYP1A activity and formation rates of this BTP in *S. trutta*. Previous reports on developmental toxicity of propranolol BTPs in different aquatic organisms (e.g. protozoans, rotifers), suggested that these compounds display similar effects than the parent compound (*32*). However, hydroxylated BTPs appear to undergo rapid phase II biotransformation, thus increasing elimination rates and reducing toxicity (*33*).

Furthermore, proposed biotransformation pathways in previous studies suggest azoxystrobin acid (AzoBP1) as the primary BTP resulting from azoxystrobin biotransformation (Fig. 2H) (*34*). However, the removal of a C_2_H_2_O group was likely responsible for the formation of AzoBP2 and AzoBP3. Interestingly, despite *L. gibbosus* displaying the highest clearance rate, azoxystrobin BTPs had higher formation rates in *S. trutta* from a site with moderate agricultural activity (Fig. 2H). Such observation could be the result of site- and species-specific factors that favored BTP production in fish from site D, or that interfered, and subsequently limited, specific enzyme-chemical interactions in fish from site F. As far as the toxic profiles of azoxystrobin BTPs, it is unclear whether these compounds could lead to similar effects than the parent compound. However, the likelihood of these BTPs undergoing phase II biotransformation, particularly glucuronidation and glutathione conjugation, is also elevated (*35, 36*), thus facilitating their excretion.

While our study indicated that clearance rates for diclofenac and terbuthylazine were low across species, one BTP was detected for each chemical in all the species that biotransformed the parent compound (Table S6). Our observations for diclofenac are in line with previous studies in fish reporting low biotransformation rates (*37*) and with other studies in salmonids indicating that hydroxylation is a major route for the formation of BTPs (*38*). However, there is a lack of information regarding the specific toxicity profiles of diclofenac BTPs in fish. In other aquatic organisms (e.g. gammarids) it appears that certain diclofenac BTPs (e.g. diclofenac methyl ester) display higher toxicity than the parent compound, likely due to higher bioaccumulation of the methylated molecule (*39*). However, such toxicity has not been associated with hydroxylated BTPs as they appear to undergo rapid glucuronidation (*38*).

Furthermore, terbuthylazine biotransformation in the evaluated fish species led to the production of TerBP1, resulting from the dealkylation of terbuthylazine, as the primary BTP. Previous studies with common carp (*Cyprinus carpio*) have indicated that the toxicity of this compound is associated with an impairment of antioxidant systems (*40*) and with a delay in growth and development of carp embryos (*41*). In such case, rapid terbuthylazine biotransformation could allow fish to cope with reactive oxygen species (ROS) and oxidative stress, and to prevent abnormal growth. Our study did not investigate the potential for bioactivation (i.e. the process of making a compound more toxic upon undergoing biotransformation) of the selected micropollutants, and assessments exploring the putative toxicity of individual BTPs would require more complex experimentation that goes beyond the scope of the present study.

Collectively, it is important to point out that BTPs represent an understudied aspect of chemical pollution (*42*), and by providing a BTP profiling for each fish species, our study advocates to generate information about BTP formation as a key consideration when estimating biotransformation and elimination kinetics of parent compounds. As analytical technologies continue to emerge and efforts to optimize bioaccumulation/biotransformation assessments are proposed (*43*), implementing practices for collecting data related to BTP formation could help in advancing knowledge across different species and environmental scenarios.

## 6. Implications for fish biodiversity in polluted waters

Our research places biotransformation as a strong indicator of species sensitivity to chemical pollution. The differential interspecific activity of important biotransformation pathways and the multiple ways in which fish species process pollutants has the potential to influence chemical body burdens (e.g. bioaccumulation) and potential adverse outcomes that may result from inhabiting polluted environments. Species that are unable to display mechanisms of defense, like biotransformation, may not be able to cope with exposure and could be at a higher risk of incurring alterations to the size and integrity of their populations. In turn, such effects could lead to severe consequences for biodiversity assemblages, as species with physiological traits that make them more resilient could overcome sensitive populations.

Therefore, we argue that conducting evaluations of biotransformation within ecological risk assessments could provide valuable information that helps in identifying species at risk and in developing appropriate measures to protect them, thus maintaining the biodiversity and ecological integrity of aquatic ecosystems.

We also provided significant observations for the need to consider the inducibility of biotransformation pathways, as this could directly determine the efficiency of species to eliminate pollutants. Historically, biotransformation studies in the laboratory have been based on the constitutive ability of organisms to biotransform chemicals, and such approach has been outlined in standardized guidelines for the estimation of bioaccumulation parameters in the context of regulation of chemicals (*22, 44*). While the conservative nature of these approaches have led to the determination of the bioaccumulation potential of hundreds of chemicals, we provided important evidence for how environmental conditions, like pollution level (i.e. exposure history), influence biotransformation activity and in consequence also bioaccumulation across species. Such consideration is imperative for research initiatives that aim to facilitate species extrapolation efforts, as the fact that even when certain species may share molecular targets for the chemicals of interest, this does not necessarily entail that the magnitude of responses and effects would be equal.

Altogether, our study clearly illustrates that the differential biotransformation responses towards micropollutant exposure could, in the long term, have important implications for fish biodiversity in contaminated waters, and that the impact of anthropogenic activity may also determine how species cope with chemical pollution. Nonetheless, we find it important to point out that a direct connection between pollution, differential biotransformation responses, and alterations to biodiversity would require long-term monitoring campaigns for the presence of different compounds as well as for the genetic diversity of fish inhabiting areas under chemical stress and their ability to display mechanisms of defense against exposure. As such, monitoring campaigns like the ones by Brodersen et al. (*11*) are essential to understand important modifications to fish biodiversity assemblages over time, and whether these modifications are linked to (in)sensitive species inhabiting polluted waters. These efforts could be also expanded to include other taxonomic groups (e.g. aquatic insects and crustaceans) that can serve as indicators of pollution-driven effects in aquatic ecosystems. In times where planetary boundaries and earth’s carrying capacity for chemical entities have been surpassed, we must continue to bring light to the detrimental consequences of chemical pollution towards biodiversity if we aim to pursue high ecosystem integrity.

## Acknowledgments

The authors would like to express their gratitude to the Fish Ecology and Evolution Department at Eawag, especially to Prof. Ole Seehausen, Dr. Dario Josi, Dr. Bernhard Wegscheider, Dr. Barbara Calegari, and Dr. Conor Waldock for their assistance in field sampling activities. The authors also thank Dr. Sarah Könemann, Dr. Christoph Schür, Dr. Fabian Balk, Dr. Mariana Torres, Sven Mosimann, Melissa von Wyl, and Pascal Bucher (Utox, Eawag), for the technical support during fish collections and processing. Also Dr. Kasia Arturi, Dr. Johannes Raths, Philipp Longree, Simon Mangold, Benedikt Lauper, and Corina Meyer (Uchem, Eawag) for their guidance during chemical analyses and data processing. Lastly, special thanks go to Dr. Rosi Siber (Siam, Eawag) for sharing her expertise in geographic information systems (GIS).

## Author Contributions

**MEF** – Conceptualization, Methodology, Investigation, Data curation, Formal Analysis, Writing -original draft; **JH** – Conceptualization, Methodology, Writing – review and editing; **KS** – Conceptualization, Methodology, Supervision, Writing – review and editing.

## Data availability

Data are available upon request.

