## Supplemental Information for "Differential biotransformation ability may alter fish biodiversity in polluted waters"

### Supplementary Information

Materials and Methods

Figs. S1 to S9

Tables S1 to S6

### Materials and Methods

**Fish and tissue sampling.** A total of six surface streams were selected within the Aare catchment across the cantons of Zürich, St. Gallen, and Schwyz in Switzerland, as to target fish species that are representatives of regional biodiversity assemblages in Switzerland and central Europe. Selected sites displayed different levels of anthropogenic activity and land use, and the site selection followed an internal classification established for freshwater biodiversity monitoring activities in the Aare catchment by the Department of Fish Ecology and Evolution at Eawag. Water quality parameters (temperature, dissolved oxygen, pH, and specific conductivity) were recorded upon arrival to each sampling location. Specific geographic and physicochemical information about all collection sites is provided in Table S1.

Across all sites, six fish species were collected: brown trout (*Salmo trutta*), common barbel (*Barbus barbus*), European bullhead (*Cottus gobio*), pumpkinseed (*Lepomis gibbosus*), and chub (*Squalius cephalus*). Fish were collected via non-lethal, backpack electrofishing, and individuals were immediately placed in aerated containers with stream water. Individuals were then separated by species and by considering similarities in size. Fish were euthanized by an overdose of buffered tricaine methanesulfonate (MS-222) and morphometric parameters (length and weight) were recorded. Fish sampling and processing were in accordance with animal experimentation regulations in Switzerland (Animal Testing Permit No. BE11/2022).

Upon dissection, approximately 30 mL of ice-cold clearing buffer (Hanks' balanced solution w/o $Ca^{2+}$/$Mg^{2+}$, 10 mM HEPES, and 2 mM EDTA, pH 7.8) were used to perfuse the liver through the hepatic vein. Once cleared, livers were excised, weighed, and immediately frozen in 1.5 mL cryovials. Liver samples were transported to the laboratory in dry ice and immediately transferred to -80°C upon arrival. The number of fish collected per species and average morphometric parameters of the individuals processed are shown in Table S2.

**Isolation of liver S9 sub-cellular fractions.** Liver S9 sub-cellular fractions were isolated from frozen tissue following standardized guidelines (*1, 2*). Pooled livers were placed in 15 mL conical tubes with two volumes homogenization buffer (50 mM Tris-HCL, 150 mM KCl, 2 mM EDTA, 1 mM DTT, and 250 mM sucrose, pH 7.8), minced, and homogenized using a ULTRA-TURRAX® digital homogenizer (IKA®-Werke GmbH & Co. KG, Staufen, DE). The resulting homogenates were centrifuged at 13,000 g for 25 min and at 4 °C (UniCn MR, Herolab GmbH, Wiesloch, DE). The resulting supernatants (S9 fractions) were then carefully collected and aliquoted in 0.5 mL cryogenic tubes. The aliquots were transferred to a -80 °C freezer until enzyme activity and substrate depletion assays were performed. Lastly, a small volume of each S9 fraction was used for total protein quantification via a modified fluorescamine method (*3*). Briefly, 25 µL of diluted S9 fraction were placed in 96-microwell plates along with 125 µL of ice-cold homogenization buffer, followed by the addition of 50 µL of a 0.3 mg/mL fluorescamine solution in acetonitrile. After a 5 min incubation at 25°C, plates were read at excitation/emission wavelengths of 360/460 nm. Protein concentrations were determined based on a calibration curve using bovine serum albumin (BSA) as standard.

**Biotransformation enzyme activity.** Michaelis-Menten kinetic parameters, including substrate concentrations that led to half of maximum reaction rates (*Km*) and maximum reaction rates (*Vmax*), were estimated by measuring the activity of different phase I and II biotransformation enzyme as a function of different substrate concentrations. Broad concentration ranges were established to obtain insight into enzyme activity kinetics under different substrate availability. These assessments included three cytochrome P450 isoforms (phase I): CYP1A, CYP2B, and CYP3A (Fig. S1), and two conjugation enzymes (phase II): glutathione S-transferase (GST) and UDP-glucuronosyltransferase (UGT; Fig. S2). The selection of enzyme substrates and bioassay design to evaluate enzymatic activity followed previously reported protocols (*4-7*).

Briefly, CYP1A activity kinetics was measured via the ethoxyresorufin-*O*-deethylase (EROD) bioassay, consisting on 0.5 mg/mL S9 protein, 2 mM NADPH, 100 mM $KPO_4$ buffer (pH 7.8), and 7-ethoxyresorufin (7-ER) in concentrations ranging from 0.05 to 50 µM in a 200 µL reaction. The dealkylation of 7-ER was measured for 5 min at 25 °C and at emission/excitation wavelengths of 535/590 nm. CYP2B-like activity was evaluated for 20 min through the pentoxyresorufin-O-deethylase (PROD) assay under similar conditions as the EROD assay, but with 7-pentoxyresorufin (7-PR) as substrate in concentrations ranging from 0.05 to 50 µM. Similarly, CYP3A-like activity kinetics was measured via the benzyloxy-4-trifluoromethylcoumarin-*O*-debenzyloxylase (BFCOD) assay in 200 µL reactions consisting of 0.5 mg/mL S9 protein, 2 mM NADPH, and benzyloxy-4-trifluoromethylcoumarin (BFC) in concentrations of 0.5 to 100 µM. The activity was monitored for 20 min at emission/excitation wavelengths of 385/500 nm.

Phase II biotransformation assays were also conducted for each fish species. The enzyme activity kinetics of GST was evaluated colorimetrically for 10 min at 340 nm, following a modified protocol (*6*). These modifications consisted in adjusting reactions to 200 µL in 96-microwell plates. Reactions included 0.1 mg/mL S9 protein, 5 mM reduced glutathione (GSH), 100 mM $KPO_4$ buffer, and 1-chloro-2,3-dinitrobenzene (CDNB) in concentrations ranging from 0.05 to 10 mM. GST activity was calculated with a conjugated CDNB molar extinction coefficient of 5.03 $mM^{-1}$ (adapted for 96-well plates). Lastly, UGT activity was monitored for 30 min by measuring reductions in absorbance at 400 nm (*7*). Reactions were also adjusted for 200 µL in 96-microwell plates, and consisted in 1 mg/mL S9 protein, 25 µg/mL alamethicin, 2 mM uridine 5'-diphosphoglucuronic acid trisodium salt (UDPGA), 100 mM $KPO_4$ buffer, and *p*-nitrophenol in concentrations ranging from 0.5 to 50 µM.

**Micropollutant depletion.** Substrate depletion assays were conducted for six micropollutants: the pharmaceuticals propranolol, diclofenac, and paracetamol, and the pesticides azoxystrobin, terbuthylazine, and pirimicarb (Fig. S3). These compounds were selected based on previous regional and global monitoring studies indicating a significant presence in surface waters (*8, 9*), and, in the case of pesticides, on their currently approved status for outdoor use (*10, 11*). For each S9 pool derived from each of the species collected at the different sites, duplicate substrate depletion experiments were established following the OECD 319b test guideline (*1*). Depletion assays consisted in 1 mL reactions with 1 mg/mL S9 protein, 2 mM of NADPH and UDPGA, 5 mM of GSH, 0.1 mM PAPS, and 25 µg/mL of alamethicin at 12 ± 1 °C, considering both the physiological temperature of the collected species and the average water temperatures at the time of sampling (Table S1). A small volume from each S9 pool was also subject to heat inactivation and employed in single depletion experiments as negative controls. Once S9 fractions and co-factors were at equilibrium, reactions were started by spiking 5 µL of the test chemical using methanol as carrier (0.5% v/v final volume). The final concentration of all test chemicals was 0.05 µM, selected based on previous experimentation with liver S9 fractions (*12*), and on reports suggesting depletion experiments to be conducted at low chemical concentrations (between 0.025-0.2 µM (*13, 14*)). The selected concentration was also within relevant levels detected in the environment (*8, 9*).

For all chemicals, depletion experiments were monitored for 90 min and 100 µL aliquots were taken at six different time points. However, initial experiments indicated that, for some species, the depletion of propranolol occurred rapidly. Thus, subsequent experimentation with this chemical was modified so that its depletion was monitored for 20, 50 and 90 min, depending on the initial results for each species. In all cases, aliquots were immediately placed in 400 µL of ice-cold methanol to stop the reaction. The resulting solution was centrifuged at 5,000*g* for 5 min at 4°C, and 200 µL of the supernatant were diluted with nanopure water containing 2.5 µg/L of the corresponding deuterated internal standard for each test chemical. The resulting solutions (20 mL) were placed in ND20 headspace crimp glass vials (BGB Analytik AG, CH) and transferred to an automated online solid phase extraction (SPE) system prior to instrumental analyses.

**Solid phase extraction and instrumental analysis.** Samples were loaded by dual injection into SPE cartridges with a flow rate of 2 mL/min using an aqueous ammonium acetate solution (2 mM). The cartridges were filled with approximately 18 mg of Oasis HLB and of an ion exchanger mix consisting of X-AX and X-CW strata (Phenomenex, Brechbüler AG, CH) and Isolute ENV+ (Biotage, SE) in a 1:1:1:5 ratio. Cartridge elution occurred with methanol containing 0.1% formic acid in back flush mode at 40 mL/min. Before chromatographic separation, the eluate was mixed to initial LC conditions using 0.1% aqueous formic acid solution.

Chemical analyses followed the methods by (*15*) and (*16*), and were conducted via high-performance liquid chromatography (HPLC; Ultimate, Thermo Fisher Scientific) coupled with hybrid quadrupole Orbitrap high-resolution tandem mass spectrometry (HR-MS/MS; Q-Exactive, Thermo Fisher Scientific) with electrospray ionization (ESI) in positive mode. Chromatographic separation employed an Atlantis® T3 column (5µm, 3.0 x 150 mm) and a mobile phase gradient of A: nanopure water and B: HPLC-grade methanol, both with 0.1% formic acid. Full scan acquisition was performed for a 100-1000 m/*z* range and a resolution of 140,000 at m/*z* 200. Target chemical quantification was conducted in Freestyle 1.7 and TraceFinder 5.1 (Thermo Fisher Scientific) and was based on internal standard calibration. The identification of biotransformation products (BTPs) was based on a suspect list (Table S3) compiled from literature (*17-19*) complemented by predictive models (BioTransformer 3.0) and metabolic logic. Identification was based on exact mass, isotope pattern and MS2 spectra. In absence of reference standards, a level of identification confidence was defined according to (*20*). The suspect screening approach used Compound Discoverer 3.3 (Thermo

Fisher Scientific; Table S3) and focused on the species and test chemicals for which depletion was significant in comparison to negative controls (Fig. S4-S9). Formation rates of the identified BTPs was conducted semi-quantitatively by using the parent compound's calibration curves followed by a plotting of the concentrations against time (h); formation rates are provided in table S6.

**Table S1.** Fish sampling locations across three different cantons in Switzerland. Sites are organized from low (A) to high (F) anthropogenic influence.

| Site ID | Stream | Canton | Coordinates | | Elevation (m)* | Anthropogenic influence | Water temperature (°C) | pH | Dissolved oxgen (mg $L^{-1}$) | Specific conductivity (µS $cm^{-1}$) | Species collected[¥] |
|---|---|---|---|---|---|---|---|---|---|---|---|
| A | Wängibach | St. Gallen | 47.209692 | 9.0854037 | 928 | Low | 11.3 | 7.7 | 10.0 | 328 | *S.trutta* |
| B | Sihl | Zürich | 47.334673 | 8.5217868 | 427 | Moderate | 14.8 | 8.4 | 10.3 | 350 | *S. trutta, B. barbus* |
| C | Chli Aa | Schwyz | 47.197968 | 8.859163 | 405 | Moderate | 12.9 | 8.2 | 10.3 | 442 | *S. cephalus* |
| D | Rufibach | St. Gallen | 47.180451 | 9.0306787 | 416 | Moderate | 16.4 | 7.4 | 10.2 | 498 | *S. trutta, C. gobio* |
| E | Jona | St. Gallen | 47.234326 | 8.837135 | 412 | High | 15.4 | 8.1 | 9.7 | 543 | *S. trutta* |
| F | Hoggibach | Schwyz | 47.175852 | 8.989479 | 412 | High | 13.0 | 7.7 | 9.0 | 480 | *S. trutta, L. gibbosus, S. cephalus* |

[¥]Estimated elevation from the Google Earth satellite imagery and geospatial platform. **Salmo trutta* (brown trout), *Barbus barbus* (common barbel), *Cottus gobio* (European bullhead), *Lepomis gibbosus* (pumpkinseed), *Squalius cephalus* (chub)

**Table S2.** Average morphometric parameters and Fulton's condition factor (K) of fish used in the preparation of liver S9 sub-cellular fractions, as well as total protein concentrations of each S9 pool prepared for the different fish species.

| Site ID | Species | S9 pool | No. Individuals | Length (cm) | Weight (g) | K | Fractional liver weight (g) | Protein concentration (mg mL$^{-1}$) |
|---|---|---|---|---|---|---|---|---|
| A | *Salmo trutta* | I | 4 | 15.0 ± 2.0 | 34.3 ± 14.2 | 1.0 ± 0.1 | 0.8 ± 0.6 | 18.8 |
| | | II | 4 | 15.4 ± 1.6 | 34.5 ± 9.4 | 0.9 ± 0.1 | 0.7 ± 0.4 | 15.6 |
| B | *Salmo trutta* | I | 3 | 8.0 ± 0.5 | 4.3 ± 0.6 | 0.8 ± 0.1 | 2.6 ± 1.0 | 5.3 |
| | | II | 2 | 16.3 ± 8.1 | 58.5 ± 70.0 | 0.9 ± 0.2 | 1.1 ± 0.4 | 14.5 |
| | *Barbus barbus* | I | 4 | 13.0 ± 0.7 | 17.3 ± 3.1 | 0.8 ± 0.0 | 1.1 ± 0.4 | 7.7 |
| C | *Squalius cephalus* | I | 3 | 16.3 ± 1.3 | 38.1 ± 13.0 | 0.8 ± 0.1 | 1.4 ± 0.4 | 24.5 |
| | | II | 3 | 16.0 ± 4.1 | 42.4 ± 29.5 | 0.9 ± 0.1 | 1.4 ± 0.1 | 27.6 |
| D | *Salmo trutta* | I | 3 | 13.6 ± 1.9 | 25.3 ± 12 | 0.9 ± 0.1 | 0.8 ± 0.2 | 17.2 |
| | | II | 4 | 9.8 ± 0.5 | 8.8 ± 1.9 | 0.9 ± 0.1 | 0.8 ± 0.2 | 5.1 |
| | *Cottus gobio* | I | 3 | 11.0 ± 1.3 | 18.7 ± 7.1 | 1.3 ± 0.1 | 2.5 ± 1.1 | 12.7 |
| | | II | 2 | 11.8 ± 1.1 | 22.5 ± 7.8 | 1.4 ± 0.1 | 2.0 ± 0.7 | 8.8 |
| E | *Salmo trutta* | I | 5 | 11.5 ± 0.6 | 15.8 ± 2.2 | 1.0 ± 0.1 | 0.9 ± 0.2 | 8.9 |
| | | II | 5 | 11.5 ± 0.4 | 15.2 ± 2.6 | 1.0 ± 0.1 | 0.9 ± 0.3 | 8.4 |
| F | *Salmo trutta* | I | 3 | 13.7 ± 1.5 | 27.9 ± 8.3 | 1.1 ± 0.0 | 1.0 ± 0.1 | 18.6 |
| | | II | 3 | 14.2 ± 1.2 | 32.1 ± 5.4 | 1.1 ± 0.1 | 1.1 ± 0.1 | 20.6 |
| | *Lepomis gibbosus* | I | 3 | 12.0 ± 0.9 | 32.8 ± 6.7 | 1.9 ± 0.0 | 1.8 ± 0.2 | 28.7 |
| | | II | 3 | 12.0 ± 0.9 | 32.9 ± 7.8 | 1.9 ± 0.0 | 1.6 ± 0.1 | 23.5 |
| | *Squalius cephalus* | I | 3 | 15.3 ± 1.3 | 33.3 ± 10.9 | 0.9 ± 0.1 | 0.8 ± 0.1 | 16.2 |
| | | II | 4 | 13.5 ± 2.4 | 26.1 ± 18.6 | 0.9 ± 0.1 | 0.9 ± 0.2 | 19.1 |

**Table S3.** List of biotransformation products (BTPs) resulting from the hepatic biotransformation of test micropollutants in liver S9 sub-cellular fractions isolated from five different fish species. The list includes the compounds that were identified following a suspect screening approach based on the reports by Rösch et al. (*18*), Jeon and Hollender (*17*) and Fu et al. (*19*). Confidence levels for BTP identification followed the approach reported by Schymanski et al. (*20*).

| Compound | Molecular Structure | Formula | Exact mass $[M+H]^+$ | RT (min) | Identification confidence level | CE [eV] | MS/MS confirmatory ions |
|---|---|---|---|---|---|---|---|
| Propranolol | | $C_{16}H_{21}NO_2$ | 260.1645 | 15.1 | 1 | 30 | 260.1649<br>183.0809<br>116.1072 |
| Propranolol-N-hydroxyl (ProBP1) | | $C_{16}H_{21}NO_3$ | 276.1597 | 12.3 | 3 | 30 | 220.4583<br>171.0806<br>74.0601 |
| 4-Hydroxypropranolol (ProBP2) | | $C_{16}H_{21}NO_3$ | 276.1597 | 12.9 | 3 | 30 | 173.0595<br>72.0808<br>58.0656 |
| Hydroxypropranolol (ProBP3) | | $C_{16}H_{21}NO_3$ | 276.1597 | 16.4 | 3 | 30 | |
| Propranolol-desisopropyl (ProBP4) | | $C_{13}H_{15}NO_2$ | 218.1183 | 14.8 | 3 | 30 | 200.2015<br>88.0759<br>70.0654 |
| Diclofenac | | $C_{14}H_{11}Cl_2NO_2$ | 296.0241 | 21.3 | 1 | 30 | 215.0498<br>214.0419 |
| Diclofenac-hydroxyl (DicBP1) | | $C_{14}H_{11}Cl_2NO_3$ | 312.0188 | 19.7 | 3 | 30 | 294.1784<br>266.1832<br>168.1102 |
| Azoxystrobin | | $C_{22}H_{17}N_3O_5$ | 404.1248 | 19.4 | 1 | 30 | 372.0983<br>344.1037<br>172.0394 |
| Azoxystrobin acid (AzoBP1) | | $C_{21}H_{15}N_3O_5$ | 390.1084 | 18.8 | 1 | 30 | 372.0981<br>302.0919<br>172.039 |
| AzoBP2 | | $C_{20}H_{15}N_3O_4$ | 362.1135 | 18.6 | 3 | 20 | 362.1135<br>302.0925<br>330.0874 |
| AzoBP3 | | $C_{20}H_{15}N_3O_4$ | 362.1135 | 19.7 | 3 | 20 | 362.1135<br>302.0925<br>330.0874 |
| Terbuthylazine | | $C_9H_{16}N_5Cl$ | 230.1167 | 20.5 | 1 | 30 | 174.0541<br>104.001<br>68.0245 |
| Terbuthylazine-desethyl (TerBP1) | | $C_7H_{12}N_5Cl$ | 202.0854 | 18.3 | 3 | 30 | 146.0229<br>79.0059<br>57.0704 |

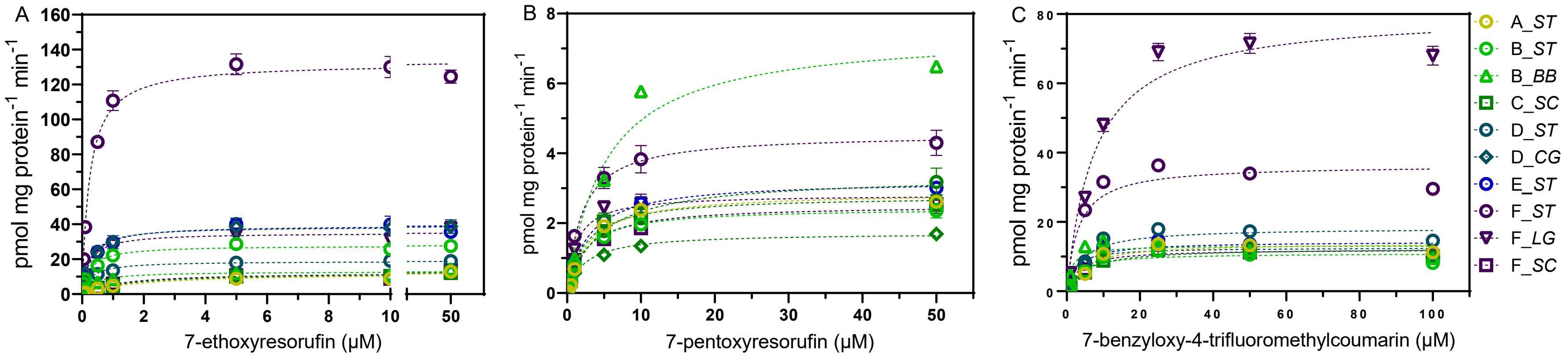


**Figure S1.** Michaelis-Menten kinetics of (A) CYP1A, (B) CYP2B-like, (C) CYP3A4-like activities in S9 sub-cellular fractions isolated from the livers of five fish species inhabiting streams with different anthropogenic influence. Points on the graphs correspond to mean enzymatic activity ± SEM of two S9 pools (one for *B. barbus*) for each species with two technical replicates. The transition from light to dark colors correspond to increasing levels of anthropogenic influence.

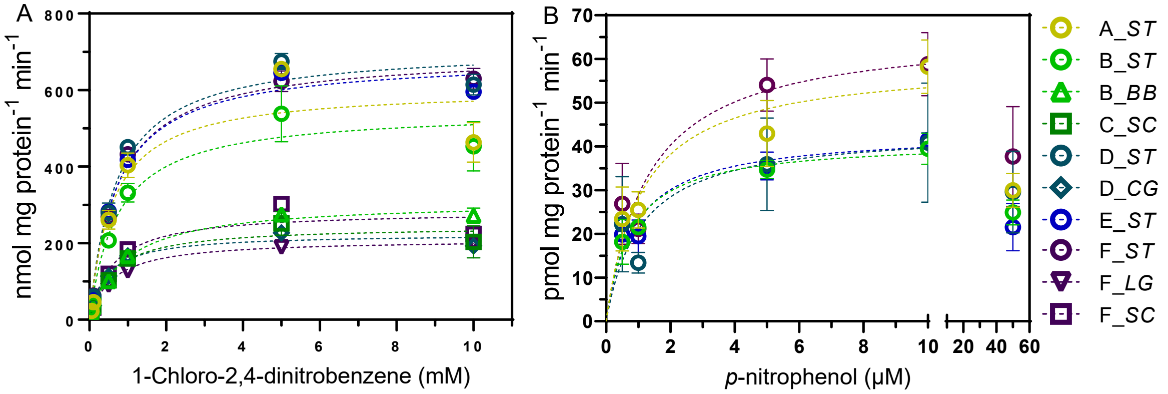


**Figure S2.** Michaelis-Menten kinetics of (A) glutathione S-transferase and (B) UDP-glucuronosyltransferase activities in S9 sub-cellular fractions isolated from the livers of five fish species inhabiting streams with different anthropogenic influence. Points on the graphs correspond to mean enzymatic activity ± SEM of two S9 pools (one for *B. barbus*) for each species with two technical replicates. The transition from light to dark colors correspond to increasing levels of anthropogenic influence.

**Table S4.** Michaelis-Menten kinetic parameters of phase I and II biotransformation enzymes in S9 sub-cellular fractions isolated from the livers of five fish species inhabiting streams with different anthropogenic influence. Values correspond to estimated substrate concentrations that resulted in half of the maximum reaction rate (*Km*; µM), maximum reaction rates (*Vmax*; pmol • mg protein$^{-1}$ min$^{-1}$), and $R^2$ derived from lines of best fit shown in Fig. 1.

| Site ID | Site_Species | CYP1A | | | CYP2B-like | | | CYP3A4-like | | | Glutathione S-transferase* | | | UDP-glucuronosyl transferase | | |
|---|---|---|---|---|---|---|---|---|---|---|---|---|---|---|---|---|
| | | ***Km*** | $V_{max}$ | $R^2$ | ***Km*** | $V_{max}$ | $R^2$ | ***Km*** | $V_{max}$ | $R^2$ | ***Km*** | $V_{max}$ | $R^2$ | ***Km*** | $V_{max}$ | $R^2$ |
| A | *Salmo trutta* | 1.8 | 12.1 | 0.85 | 2.9 | 2.9 | 0.98 | 4.8 | 13.9 | 0.49 | 0.6 | 605.0 | 0.87 | 1.1 | 59.4 | 0.55 |
| B | *Salmo trutta* | 0.3 | 27.8 | 0.96 | 2.4 | 2.4 | 0.96 | 3.3 | 11.0 | 0.79 | 0.7 | 546.7 | 0.85 | 0.8 | 41.3 | 0.69 |
| B | *Barbus barbus* | 0.4 | 12.9 | 0.93 | 5.1 | 7.5 | 0.98 | 1.3 | 13.2 | 0.75 | 0.9 | 309.6 | 0.99 | -- | -- | -- |
| C | *Squalius cephalus* | 1.5 | 12.4 | 0.87 | 2.4 | 2.8 | 0.92 | 5.2 | 12.4 | 0.78 | 0.6 | 244.7 | 0.79 | -- | -- | -- |
| D | *Salmo trutta* | 0.3 | 18.9 | 0.93 | 4.4 | 3.4 | 0.93 | 4.1 | 18.3 | 0.83 | 0.7 | 711.3 | 0.97 | 1.1 | 44.0 | 0.20 |
| D | *Cottus gobio* | 0.3 | 38.7 | 0.77 | 2.5 | 1.7 | 0.97 | 2.0 | 12.6 | 0.71 | 0.4 | 224.0 | 0.95 | 3.2 | 80.2 | 0.22 |
| E | *Salmo trutta* | 0.3 | 39.3 | 0.85 | 2.8 | 3.2 | 0.98 | 3.2 | 14.3 | 0.85 | 0.7 | 683.8 | 0.98 | 0.9 | 43.2 | 0.74 |
| F | *Salmo trutta* | 0.2 | 132.4 | 0.96 | 2.0 | 4.6 | 0.94 | 3.2 | 36.4 | 0.84 | 0.7 | 696.9 | 0.98 | 1.3 | 66.5 | 0.58 |
| F | *Lepomis gibbosus* | 0.2 | 35.1 | 0.93 | 1.3 | 2.8 | 0.93 | 7.6 | 80.4 | 0.95 | 0.6 | 209.9 | 0.98 | -- | -- | -- |
| F | *Squalius cephalus* | 1.3 | 12.6 | 0.90 | 2.6 | 2.5 | 0.96 | 3.4 | 12.1 | 0.54 | 0.6 | 284.0 | 0.90 | -- | -- | -- |

**Km* given in mM of substrate; *Vmax* given as nmol • mg protein$^{-1}$ min$^{-1}$.

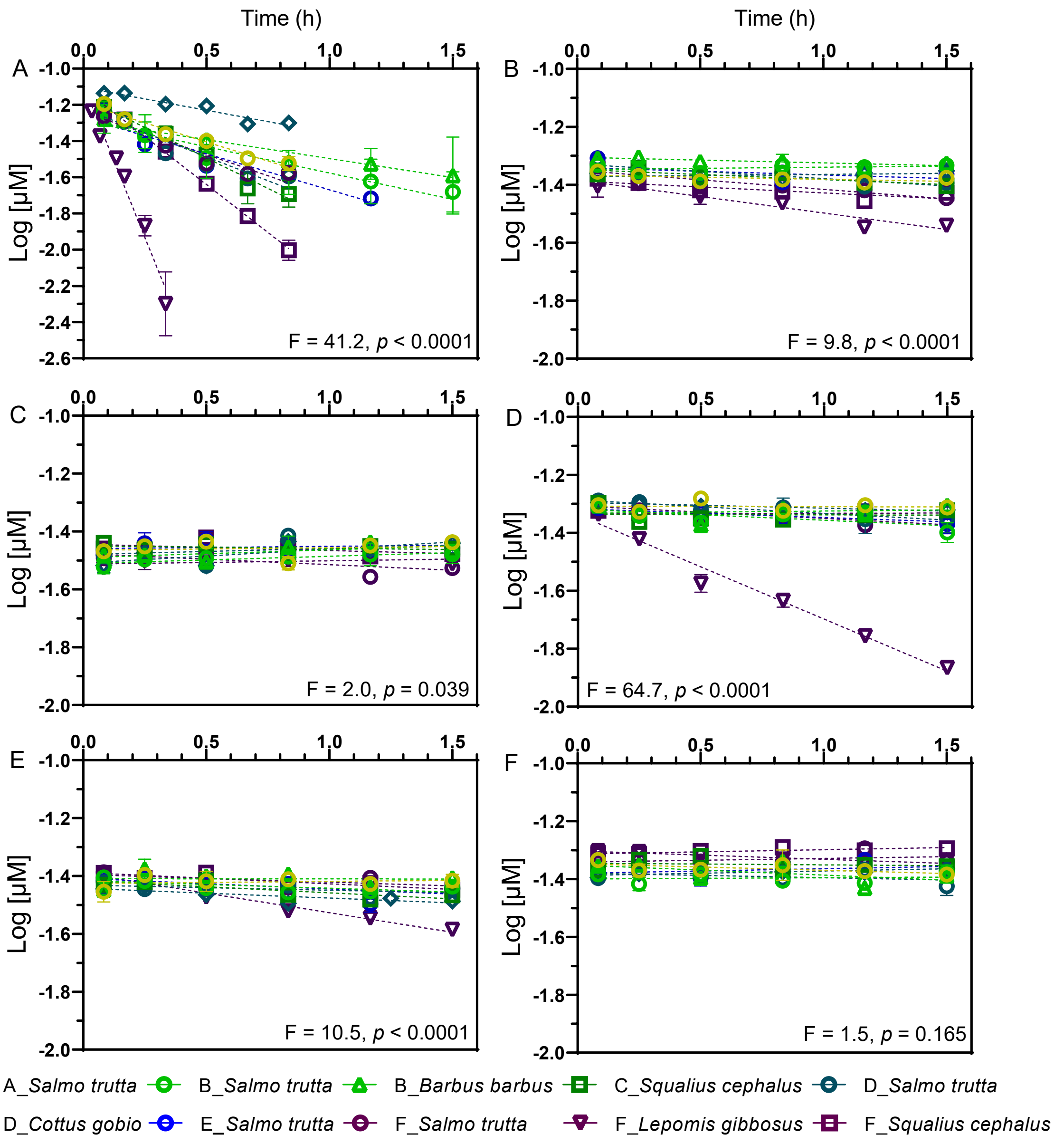


**Figure S3.** Depletion of (A) propranolol, (B) diclofenac, and (C) paracetamol, (D) azoxystrobin, (E) terbuthylazine, and (F) pirimicarb by liver S9 sub-cellular fractions isolated from five fish species inhabiting streams with different anthropogenic influence. Data points correspond to the mean $Log_{10}$ concentrations (µM) ± SEM of two independent experiments with each S9 pool (n = 4 for most species and n = 2 for *B. barbus*). Each graph indicates whether the slopes resulting from linear regression analyses are different from each other, when considering $p < 0.05$ as significant. The transition from light to dark colors correspond to increasing levels of anthropogenic influence.

**Table S5.** *In vitro* hepatic clearance rates ($CL_{IN\ VITRO}$) of the six test micropollutants by S9 sub-cellular fractions isolated from the livers of five fish species inhabiting streams with different anthropogenic influence. $R^2$ values were obtained from linear regression analyses performed for each species-chemical combination (Fig. 2). Missing values indicate no chemical depletion relative to negative controls (Figs. S1 – S6).

| Site ID | Species | Propranolol | | Diclofenac | | Paracetamol | | Azoxystrobin | | Terbuthylazine | | Pirimicarb | |
|---|---|---|---|---|---|---|---|---|---|---|---|---|---|
| | | $CL_{INVITRO}$ | $R^2$ | $CL_{INVITRO}$ | $R^2$ | $CL_{INVITRO}$ | $R^2$ | $CL_{INVITRO}$ | $R^2$ | $CL_{INVITRO}$ | $R^2$ | $CL_{INVITRO}$ | $R^2$ |
| A | *Salmo trutta* | 0.97 | 0.96 | -- | -- | -- | -- | -- | -- | -- | -- | -- | -- |
| B | *Salmo trutta* | 0.67 | 0.91 | -- | -- | -- | -- | -- | -- | 0.06 | 0.48 | -- | -- |
| B | *Barbus barbus* | 0.49 | 0.95 | 0.04 | 0.62 | -- | -- | -- | -- | -- | -- | -- | -- |
| C | *Squalius cephalus* | 1.55 | 0.98 | 0.08 | 0.70 | -- | -- | -- | -- | 0.10 | 0.68 | -- | -- |
| D | *Salmo trutta* | 1.22 | 0.88 | 0.12 | 0.84 | -- | -- | 0.12 | 0.86 | -- | -- | -- | -- |
| D | *Cottus gobio* | 0.57 | 0.92 | -- | -- | -- | -- | 0.05 | 0.94 | 0.08 | 0.66 | -- | -- |
| E | *Salmo trutta* | 0.92 | 0.91 | -- | -- | -- | -- | 0.07 | 0.73 | 0.09 | 0.47 | -- | -- |
| F | *Salmo trutta* | 1.08 | 0.91 | 0.15 | 0.76 | 0.09 | 0.45 | 0.09 | 0.76 | 0.06 | 0.59 | -- | -- |
| F | *Lepomis gibbosus* | 7.76 | 0.97 | 0.26 | 0.92 | -- | -- | 0.83 | 0.98 | 0.31 | 0.97 | 0.07 | 0.90 |
| F | *Squalius cephalus* | 2.44 | 0.99 | 0.15 | 0.95 | -- | -- | -- | -- | 0.09 | 0.72 | -- | -- |

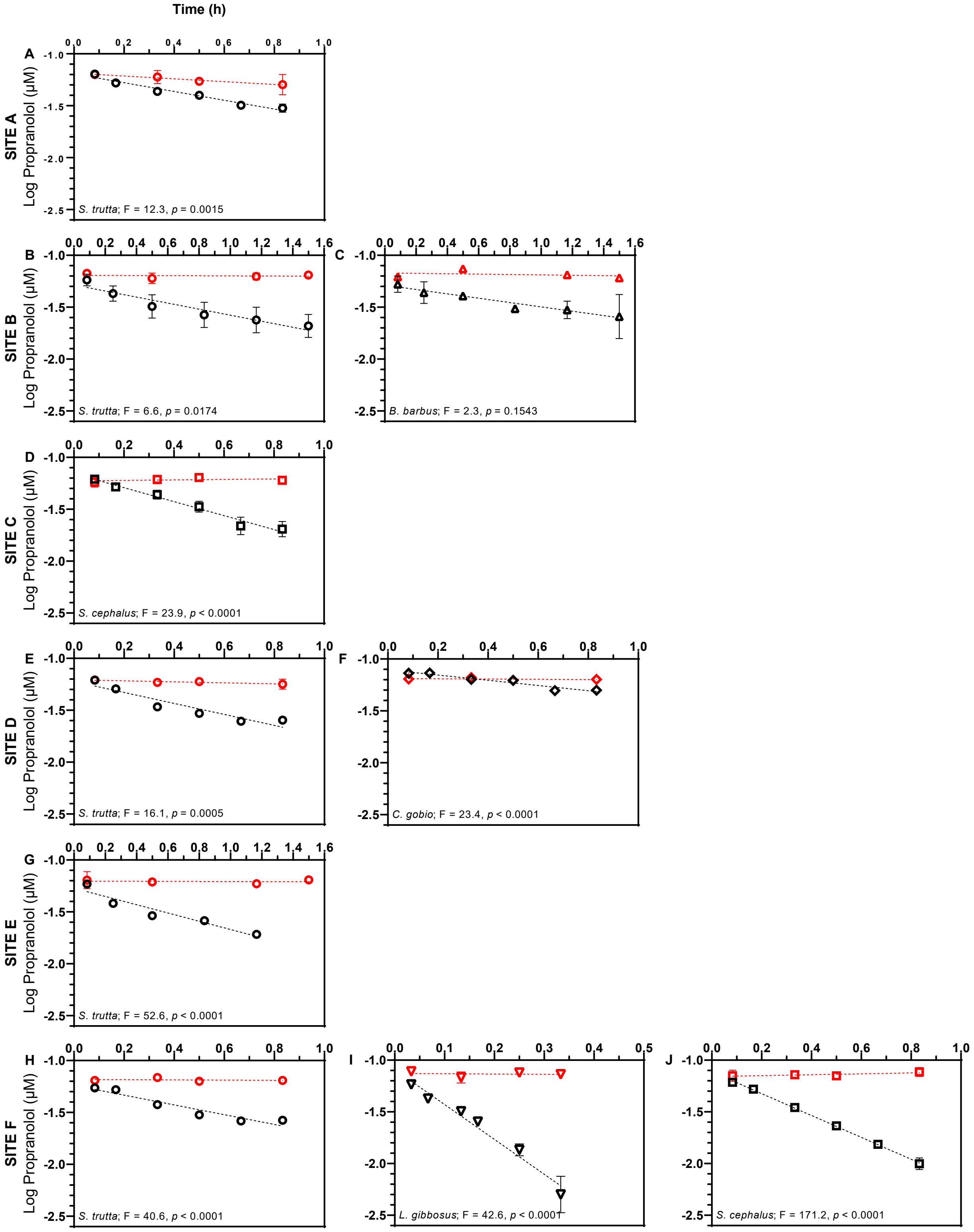


**Figure S4.** Depletion of propranolol by active (black) and heat-inactivated (red) liver S9 sub-cellular fractions isolated from five fish species inhabiting streams with different anthropogenic influence. Panels are organized as to show individual species collected from the different sites. Data points correspond to the mean $Log_{10}$ concentrations (µM) ± SEM of two independent experiments (one for heat-inactivated S9) with each S9 pool (n = 4 for most species and n = 2 for *B. barbus*). Each graph indicates whether the slopes resulting from linear regression analyses are different from each other, when considering $p < 0.05$ as significant.

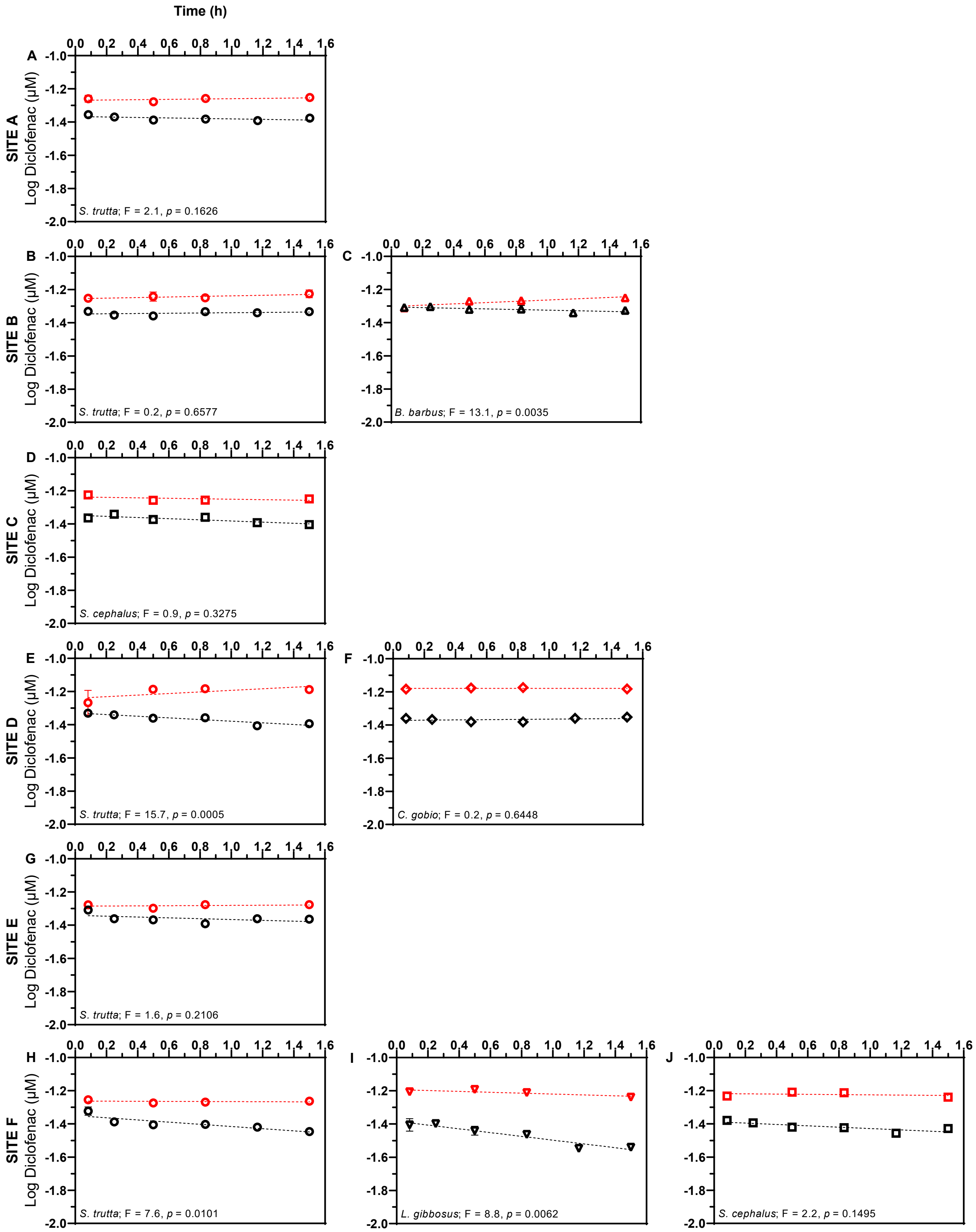


**Figure S5.** Depletion of diclofenac by active (black) and heat-inactivated (red) liver S9 sub-cellular fractions isolated from five fish species inhabiting streams with different anthropogenic influence. Panels are organized as to show individual species collected from the different sites. Data points correspond to the mean $Log_{10}$ concentrations (µM) ± SEM of two independent experiments (one for heat-inactivated S9) with each S9 pool (n = 4 for most species and n = 2 for *B. barbus*). Each graph indicates whether the slopes resulting from linear regression analyses are different from each other, when considering p < 0.05 as significant.

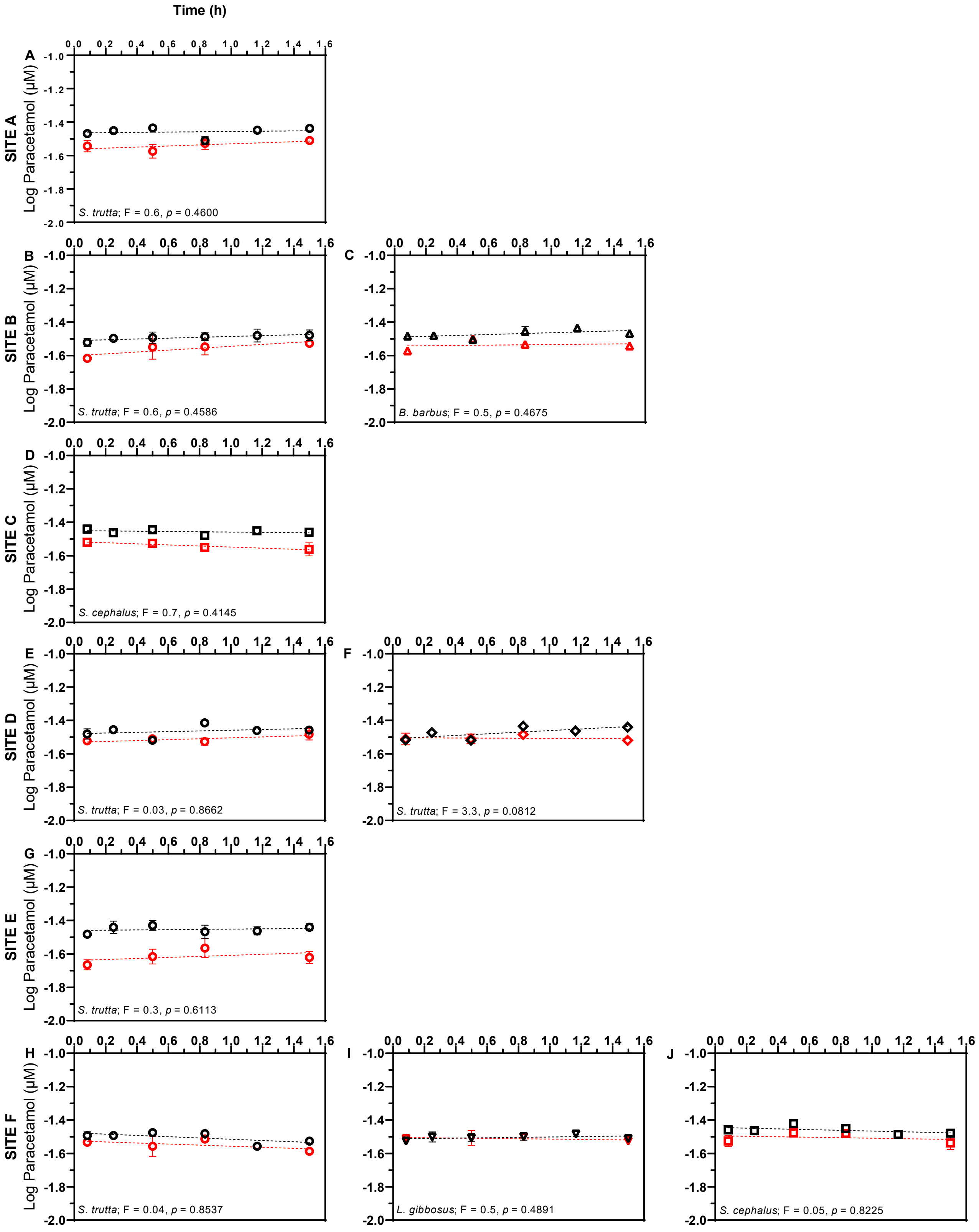


**Figure S6.** Depletion of paracetamol by active (black) and heat-inactivated (red) liver S9 sub-cellular fractions isolated from five fish species inhabiting streams with different anthropogenic influence. Panels are organized as to show individual species collected from the different sites. Data points correspond to the mean $Log_{10}$ concentrations (µM) ± SEM of two independent experiments (one for heat-inactivated S9) with each S9 pool (n = 4 for most species and n = 2 for *B. barbus*). Each graph indicates whether the slopes resulting from linear regression analyses are different from each other, when considering p < 0.05 as significant.

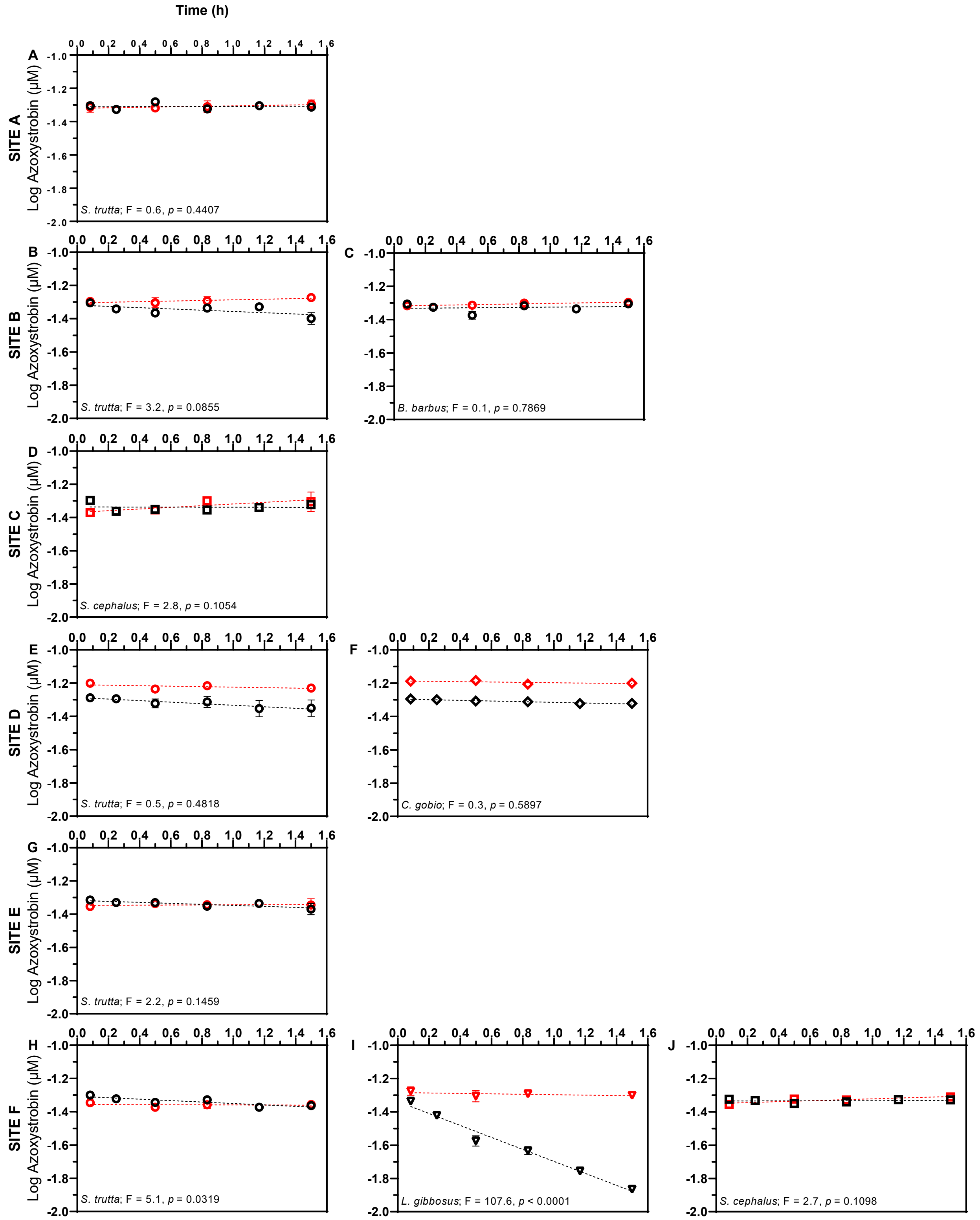


**Figure S7.** Depletion of azoxystrobin by active (black) and heat-inactivated (red) liver S9 sub-cellular fractions isolated from five fish species inhabiting streams with different anthropogenic influence. Panels are organized as to show individual species collected from the different sites. Data points correspond to the mean $Log_{10}$ concentrations (µM) ± SEM of two independent experiments (one for heat-inactivated S9) with each S9 pool (n = 4 for most species and n = 2 for *B. barbus*). Each graph indicates whether the slopes resulting from linear regression analyses are different from each other, when considering $p < 0.05$ as significant.

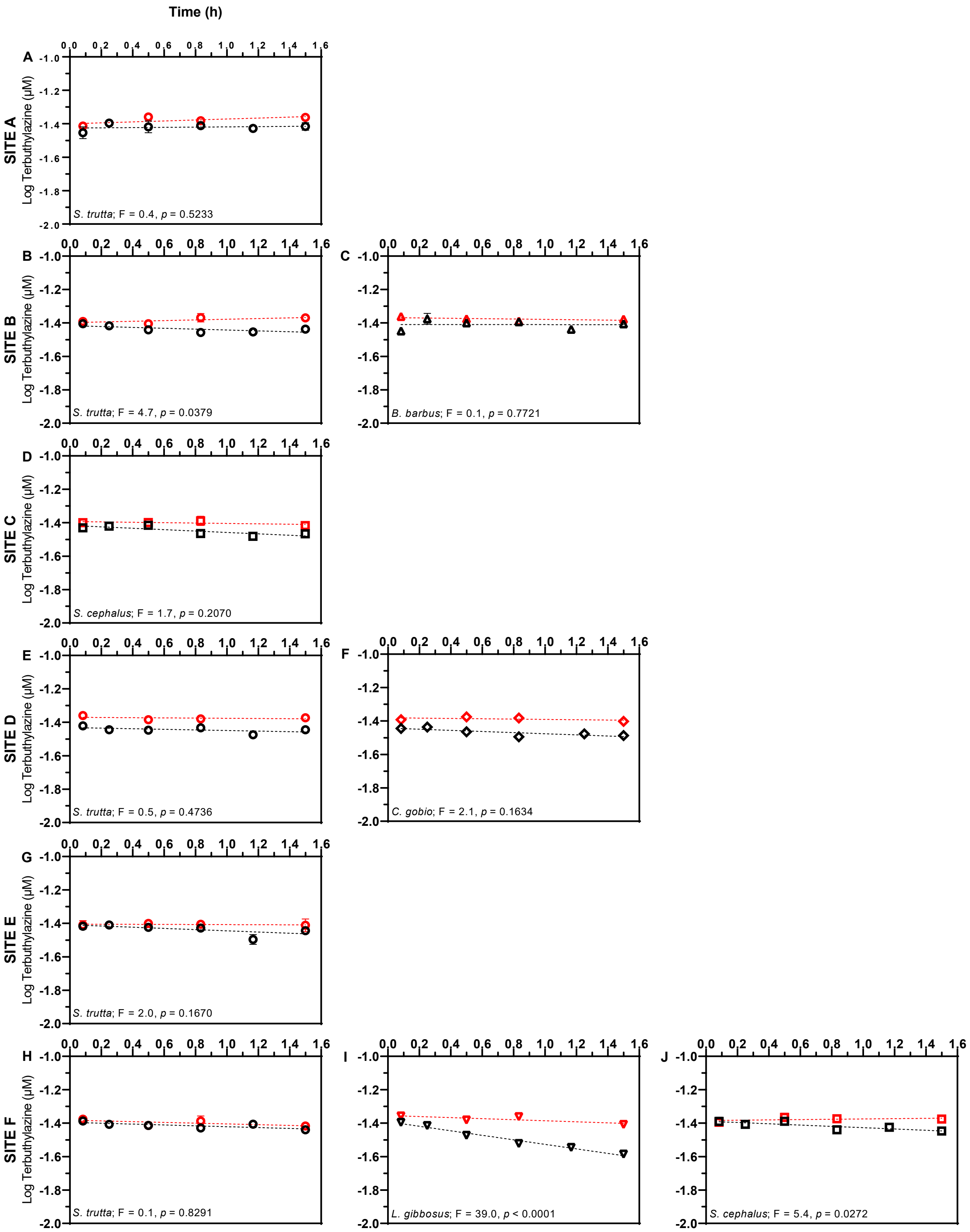


**Figure S8.** Depletion of terbuthlyazine by active (black) and heat-inactivated (red) liver S9 sub-cellular fractions isolated from five fish species inhabiting streams with different anthropogenic influence. Panels are organized as to show individual species collected from the different sites. Data points correspond to the mean $Log_{10}$ concentrations (µM) ± SEM of two independent experiments (one for heat-inactivated S9) with each S9 pool (n = 4 for most species and n = 2 for *B. barbus*). Each graph indicates whether the slopes resulting from linear regression analyses are different from each other, when considering p < 0.05 as significant.

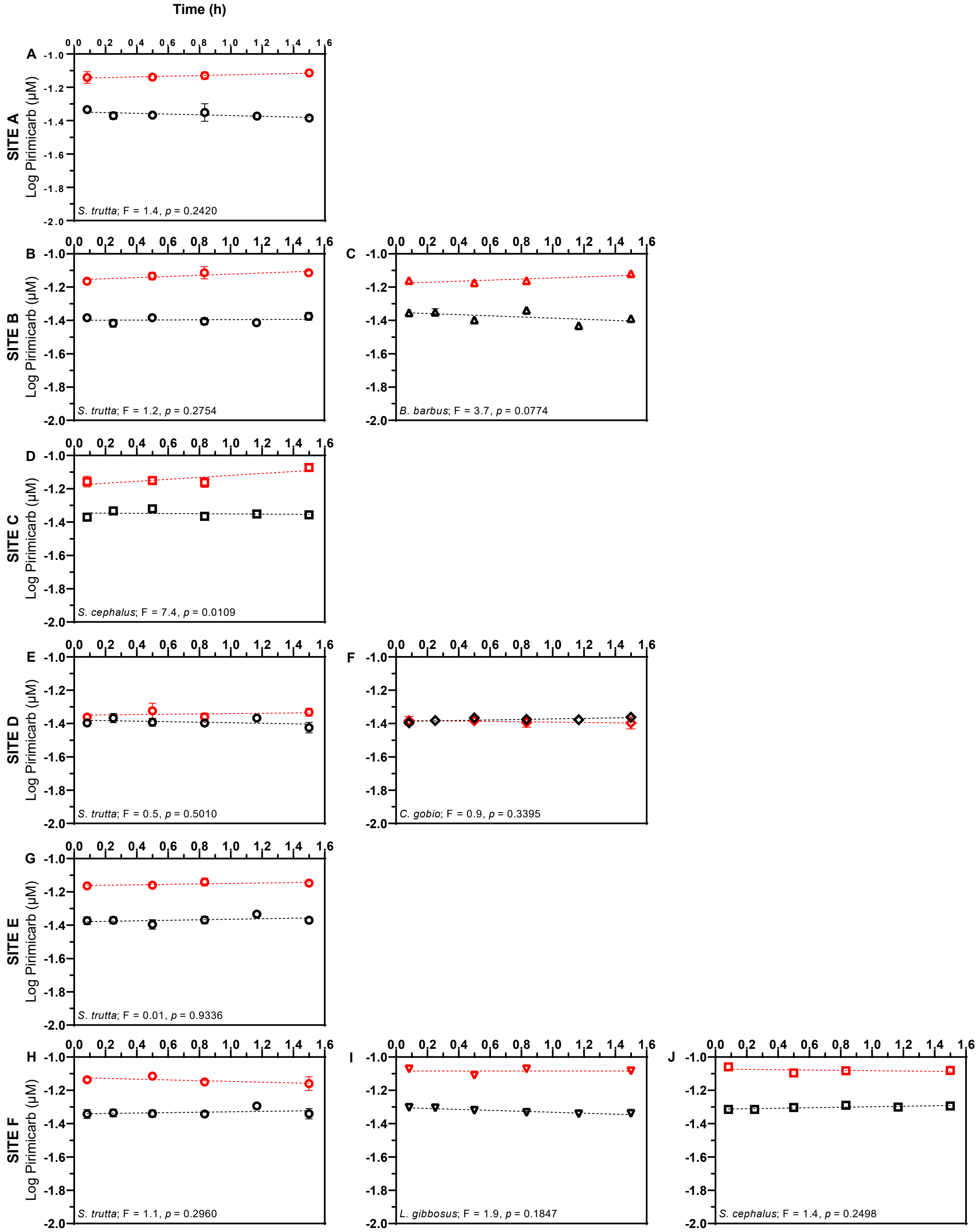


**Figure S9.** Depletion of pirimicarb by active (black) and heat-inactivated (red) liver S9 sub-cellular fractions isolated from five fish species inhabiting streams with different anthropogenic influence. Panels are organized as to show individual species collected from the different sites. Data points correspond to the mean $Log_{10}$ concentrations (µM) ± SEM of two independent experiments (one for heat-inactivated S9) with each S9 pool (n = 4 for most species and n = 2 for *B. barbus*). Each graph indicates whether the slopes resulting from linear regression analyses are different from each other, when considering $p < 0.05$ as significant.

**Table S6.** Time-dependent formation (pmoles/h) of biotransformation products (BTPs) resulting from the hepatic depletion of PRO, DIC, AZO, and TER, and $R^2$ of the lines of best fit when plotting concentrations against time. Missing values indicate that the BTP was not identified or that its formation did not display a time-dependent increase.

| Site ID | Species | ProBP1 | | ProBP2 | | ProBP3 | | ProBP4 | |
|---|---|---|---|---|---|---|---|---|---|
| | | Rate | $R^2$ | Rate | $R^2$ | Rate | $R^2$ | Rate | $R^2$ |
| A | *Salmo trutta* | 0.74 | 0.89 | 3.18 | 0.85 | 15.98 | 0.84 | 4.73 | 0.70 |
| B | *Salmo trutta* | -- | -- | 0.74 | 0.27 | 4.85 | 0.33 | -- | -- |
| B | *Barbus barbus* | 0.32 | 0.44 | 0.29 | 0.21 | 0.49 | 0.44 | -- | -- |
| C | *Squalius cephalus* | 2.45 | 0.72 | 28.31 | 0.49 | 1.49 | 0.86 | -- | -- |
| D | *Salmo trutta* | 0.69 | 0.68 | 4.41 | 0.86 | 19.36 | 0.80 | -- | -- |
| D | *Cottus gobio* | -- | -- | 0.80 | 0.60 | 0.83 | 0.87 | 0.53 | 0.78 |
| E | *Salmo trutta* | 0.37 | 0.79 | 1.96 | 0.77 | 8.12 | 0.66 | -- | -- |
| F | *Salmo trutta* | 0.67 | 0.89 | 4.09 | 0.90 | 8.92 | 0.85 | 2.22 | 0.57 |
| F | *Lepomis gibbosus* | 33.07 | 0.93 | 32.41 | 0.83 | 3.17 | 0.89 | 2.13 | 0.46 |
| F | *Squalius cephalus* | 3.32 | 0.93 | 31.12 | 0.61 | 0.94 | 0.75 | -- | -- |

| | | DicBP1 | |
|---|---|---|---|
| | | Rate | $R^2$ |
| A | *Salmo trutta* | -- | -- |
| B | *Salmo trutta* | -- | -- |
| B | *Barbus barbus* | 0.97 | 0.96 |
| C | *Squalius cephalus* | -- | -- |
| D | *Salmo trutta* | 0.18 | 0.46 |
| D | *Cottus gobio* | -- | -- |
| E | *Salmo trutta* | -- | -- |
| F | *Salmo trutta* | 0.13 | 0.71 |
| F | *Lepomis gibbosus* | 6.32 | 0.96 |
| F | *Squalius cephalus* | 0.24 | 0.83 |

| | | AzoBP1 | | AzoBP2 | | AzoBP3 | |
|---|---|---|---|---|---|---|---|
| | | Rate | $R^2$ | Rate | $R^2$ | Rate | $R^2$ |
| A | *Salmo trutta* | -- | -- | -- | -- | -- | -- |
| B | *Salmo trutta* | -- | -- | -- | -- | -- | -- |
| B | *Barbus barbus* | -- | -- | -- | -- | -- | -- |
| C | *Squalius cephalus* | -- | -- | -- | -- | -- | -- |
| D | *Salmo trutta* | 3.29 | 0.72 | 0.22 | 0.92 | 0.14 | 0.50 |
| D | *Cottus gobio* | 0.67 | 0.75 | 0.05 | 0.85 | -- | -- |
| E | *Salmo trutta* | 1.32 | 0.85 | 0.11 | 0.73 | -- | -- |
| F | *Salmo trutta* | 0.82 | 0.71 | 0.06 | 0.46 | 0.14 | 0.41 |
| F | *Lepomis gibbosus* | 0.96 | 0.80 | 0.11 | 0.72 | -- | -- |
| F | *Squalius cephalus* | -- | -- | -- | -- | -- | -- |

| | | TerBP1 | |
|---|---|---|---|
| | | Rate | $R^2$ |
| A | *Salmo trutta* | -- | -- |
| B | *Salmo trutta* | 0.01 | 0.12 |
| B | *Barbus barbus* | -- | -- |
| C | *Squalius cephalus* | 0.04 | 0.75 |
| D | *Salmo trutta* | -- | -- |
| D | *Cottus gobio* | 0.20 | 0.69 |
| E | *Salmo trutta* | 0.04 | 0.81 |
| F | *Salmo trutta* | 0.16 | 0.97 |
| F | *Lepomis gibbosus* | 0.59 | 0.70 |
| F | *Squalius cephalus* | 0.02 | 0.40 |
